# Pan-cancer landscape of cancer-testis genes revealed by single-cell and spatial transcriptomics

**DOI:** 10.1101/2025.08.25.672079

**Authors:** Chunyang Fu, Shumin Yin, Linjin Li, Xiuyuan Jin, Xueying Liu, Ke Liu

## Abstract

Cancer-testis genes (CTGs) are attractive immunotherapeutic targets owing to their restricted testicular expression and aberrant activation in cancers. However, their regulatory mechanisms, spatial organization, and clinical utility remain incompletely understood. Here, we leveraged large-scale single-cell and spatial transcriptomic data to perform a comprehensive pan-cancer analysis of CTGs. We established a high-confidence pan-cancer CTG catalog and uncovered a heterogeneous epigenetic regulatory landscape in which X-linked CTGs are predominantly governed by DNA methylation, whereas autosomal CTGs are more strongly associated with chromatin regulators. Building on the observation that CTG activation is a robust pan-cancer hallmark of malignancy, we developed a computational framework that enables rapid malignant cell annotation with performance comparable to established copy number variation-based methods. Spatial transcriptomic analyses revealed that CTG expression in head and neck squamous cell carcinoma is preferentially enriched at the invasive tumor front. Clinically, we highlighted the underappreciated therapeutic potential of CTGs and prioritized *CT83* and *DCAF4L2* as promising candidate targets for T-cell receptor-engineered T-cell therapy in triple-negative breast cancer and liver cancer, respectively. Our study advances mechanistic understanding of CTG biology and provides a valuable resource for the development of CTG-based immunotherapies.

## Introduction

Cancer-testis genes (CTGs) represent a distinctive gene class characterized by their high expression in the testis, aberrant activation in various cancer types, and minimal or absent expression in most normal tissues^1,2^. Owing to the immune-privileged status of the testis, many CTG-encoded proteins—collectively termed cancer-testis antigens (CTAs)—are inherently immunogenic and have therefore attracted substantial interest as targets for cancer immunotherapy^3,4^. Beyond their immunogenicity, emerging evidence indicates that CTGs actively participate in multiple oncogenic programs, including malignant proliferation^5^, metastatic dissemination^6^, immune evasion^7,8^, and therapeutic resistance^9^.

Given this broad biological and clinical relevance, constructing a comprehensive and accurate CTG catalog is a fundamental prerequisite for both mechanistic and translational studies. Early efforts, exemplified by the CTdatabase^10^, established the first curated resource by compiling more than 200 CTGs reported in the literature. In recent years, large-scale omics-based pan-cancer analyses have substantially advanced cancer research, and similar approaches have been applied to CTGs. For example, Wang et al.^11^ expanded the CTG catalog through integrative analysis of bulk transcriptomic data from The Cancer Genome Atlas (TCGA) and the Genotype-Tissue Expression (GTEx) project, while Carter et al.^12^ identified 103 CTGs activated across multiple tumor types and demonstrated the anti-tumor efficacy of Lin28a- and Siglece-based vaccination strategies. Although these studies advanced the field, several fundamental questions remain difficult to address because of the intrinsic limitations of bulk tissue sequencing. First, the conflation of tumor-cell expression with expression from non-malignant cells hinders precise delineation of the regulatory mechanisms underlying CTG reactivation in malignant cells. Second, bulk tissue profiling obscures intratumoral heterogeneity, making it difficult to determine whether CTG activation is ubiquitous or restricted to specific malignant subpopulations. Third, the spatial organization of CTG-expressing cells and their interactions with the local tumor microenvironment have not been systematically explored.

The rapid accumulation of publicly available cancer single-cell and spatial transcriptomic data has created an unprecedented opportunity to address these limitations^13–17^. In this study, we performed a comprehensive pan-cancer analysis of CTGs using single-cell and spatial transcriptomic data from 1,326 tumor samples (**Fig. 1a**). We constructed a refined, high-quality pan-cancer CTG catalog and delineated the regulatory and spatial heterogeneity of CTG activation across tumors. We demonstrated the utility of CTGs for malignancy annotation and developed MaligCTG, a computational framework for rapid malignant cell identification with performance comparable to conventional methods. Furthermore, we highlighted the underexplored potential of CTGs in T-cell receptor-engineered T-cell (TCR-T) therapy and prioritized promising candidate targets for TCR-T therapy in triple-negative breast cancer and liver cancer.

**Fig. 1.**
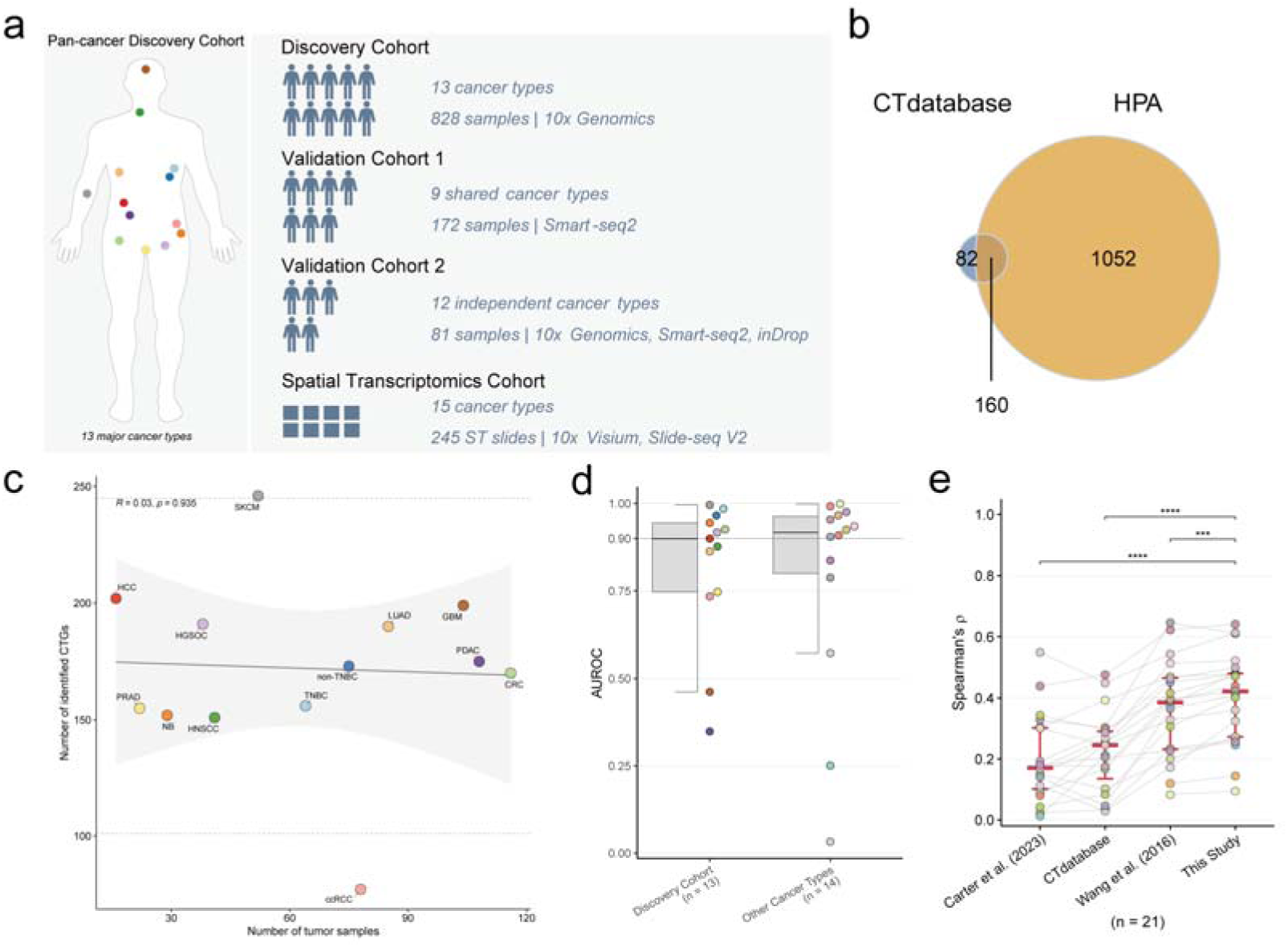
Establishment of a pan-cancer CTG catalog. **(a)** Schematic overview of the single-cell/nucleus RNA-seq cohorts and spatial transcriptomics cohort used in this study. Colors representing cancer types in the Discovery Cohort are consistent across all figures. **(b)** Venn diagram showing the overlap of CTG candidate genes from the CTdatabase and HPA. **(c)** Scatter plot illustrating the relationship between the number of tumor samples (x-axis) and the number of identified CTGs (y-axis) across cancer types. The solid grey line represents a linear regression fit, with the shaded region indicating the 95% confidence interval. Spearman correlation and *P*-value are shown. Horizontal dashed lines indicate the outlier boundaries defined by the interquartile range (IQR) method (Q1 − 1.5 × IQR and Q3 + 1.5 × IQR). **(d)** Boxplots of AUROC values quantifying the performance of aggregated CTG expression in distinguishing tumor from normal samples across cancer types in the Discovery Cohort and other TCGA cancer types. Each point represents a cancer type. **(e)** Line-connected dot plot comparing the Spearman correlations between gene-set expression and tumor purity for TCGA cancer types. The analyzed gene sets include CTGs from CTdatabase, Wang et al., Carter et al., and this study. The horizontal red bars indicate the median values, and the vertical red error bars represent the IQR. *P*-values were determined using two-sided paired Wilcoxon signed-rank tests. Each point represents a cancer type. Boxplots represent the median (center line), upper and lower quartiles (box limits), and 1.5 × IQR (whiskers). Significance: \**P* ≤ 0.05, \*\**P* ≤ 0.01, \*\*\**P* ≤ 0.001, \*\*\*\**P* ≤ 0.0001; ns, not significant.

## Results

### Establishment of a pan-cancer CTG catalog

To identify CTGs across multiple cancer types using single-cell transcriptomic data, we first assembled a Discovery Cohort comprising 828 single-cell and single-nucleus RNA-seq (scRNA-seq and snRNA-seq) samples sequenced on 10x Genomics platforms (**Fig. 1a and Supplementary Fig. 1a–c**). This cohort encompassed 13 major cancer types, including glioblastoma (GBM), breast cancer (BRCA; subtyped into triple-negative [TNBC] and non-triple-negative [non-TNBC]), colorectal cancer (CRC), head and neck squamous cell carcinoma (HNSCC), clear cell renal cell carcinoma (ccRCC), hepatocellular carcinoma (HCC), lung adenocarcinoma (LUAD), neuroblastoma (NB), high-grade serous ovarian cancer (HGSOC), pancreatic ductal adenocarcinoma (PDAC), prostate cancer (PRAD), and melanoma (SKCM). Next, we compiled a list of 1,294 unique CTG candidate genes from two sources: 242 curated CTGs from the CTdatabase and 1,212 testis-enriched protein-coding genes from the Human Protein Atlas^18^ (**Fig. 1b**). As expected, most of these genes showed peak expression exclusively in testicular germ cells among normal human cell types (**Supplementary Fig. 1d**).

We defined a gene *g* as a CTG for cancer type *C* if its expression was detected in more than 1% of malignant cells in at least half of the samples within *C*. This stringent criterion yielded 407 unique CTGs, with a median of 173 CTGs per cancer type. SKCM exhibited the highest number of CTGs, whereas ccRCC showed the lowest, consistent with its established CTG-poor phenotype^19^ (**Fig. 1c**). Notably, we did not observe a significant correlation between the number of CTGs and either sample size or the number of detected genes across cancer types (**Fig. 1c** and **Supplementary Fig. 2a**), suggesting that variation in CTG abundance reflects intrinsic tumor biology rather than merely technical artifacts. For each CTG, we counted the number of cancer types in which it was detected, and found the resulting distribution appeared to be bimodal, suggesting that most CTGs are either highly specific to a particular cancer type or broadly expressed across multiple cancer types (**Supplementary Fig. 2b**).

To assess the quality of the identified CTGs, we examined their ability to distinguish tumor from normal samples using bulk RNA-seq data from TCGA^20^ and GTEx^21^. When aggregated CTG expression was used as a classification feature, the median area under the receiver operating characteristic curve (AUROC) reached 0.90 across the 13 cancer types in the Discovery Cohort (**Fig. 1d**). Consistent with this, CTG expression was positively correlated with tumor purity in most cancer types (median Spearman’s ρ = 0.48; **Supplementary Fig. 2c–d**). Similar results were obtained even in cancer types not included in the Discovery Cohort, underscoring the generalizability of our CTG set (**Fig. 1d** and **Supplementary Fig. 2c**). Critically, our refined CTG set exhibited higher correlation coefficients with tumor purity than previously established CTG sets^10–12^ (**Fig. 1e**), supporting the value of defining CTGs using single-cell transcriptomic data.

In summary, we identified 407 CTGs across 13 human cancer types through large-scale analysis of cancer single-cell transcriptomic datasets and validated them in independent datasets.

### Methylation-dependent gating restricts X-linked CTG activation

DNA methylation has been identified as an important regulator of CTG activation for several canonical X-linked CTGs^22–24^. However, whether this regulatory relationship extends to all CTGs remains unclear. To examine this, we assessed the association between promoter DNA methylation and gene expression across TCGA samples. Autosomal CTGs showed correlations centered near zero or only weakly negative, whereas X-linked CTGs exhibited a pronounced negative correlation across multiple cancer types (**Fig. 2a** and **Supplementary Fig. 3a**), suggesting that activation of X-linked CTGs is largely governed by promoter DNA methylation. In particular, HCC exhibited lower global DNA methylation levels than other cancer types (**Fig. 2b**), which may explain why this cancer type harbored the highest number of X-linked CTGs in our Discovery Cohort (**Supplementary Fig. 3b**). To further interrogate this dependency, we analyzed bulk RNA-seq data from HCC cell lines following treatment with the DNA methyltransferase (DNMT) inhibitor 5-Aza-2′-deoxycytidine (5-Aza-CdR)^25^. Consistent with the correlative analysis, 5-Aza-CdR treatment robustly induced X-linked CTG expression, whereas autosomal CTGs remained largely unchanged (**Fig. 2c**).

**Fig. 2.**
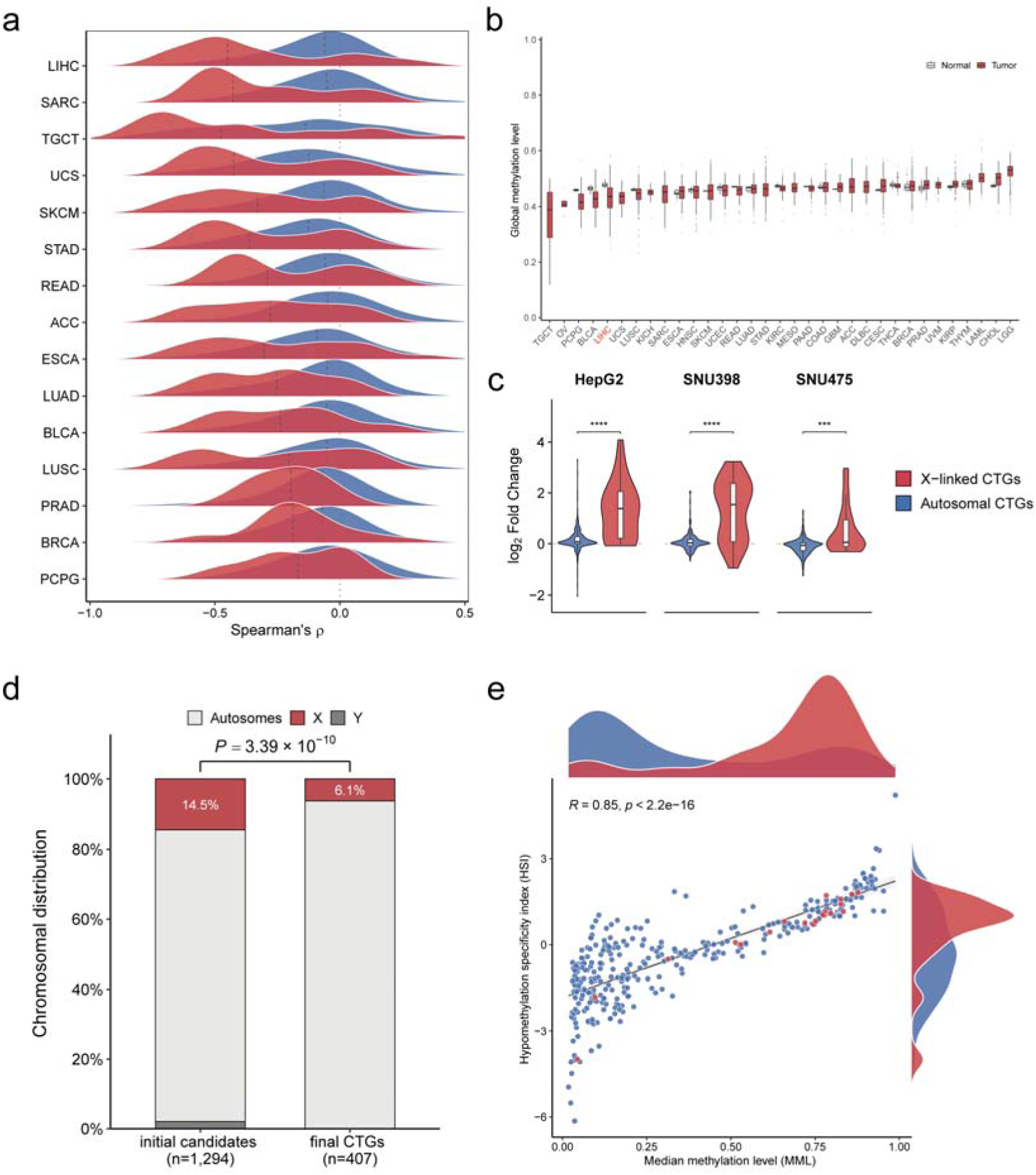
Methylation-dependent gating restricts X-linked CTG activation. **(a)** Ridge plots illustrating the distribution of Spearman correlation coefficients between promoter methylation and gene expression for autosomal CTGs and X-linked CTGs. Vertical black lines indicate the median correlation. The plot displays the top 15 TCGA cancer types showing the largest difference in correlation medians between the two groups. **(b)** Distribution of global methylation levels across cancer types from the TCGA project. Cancer types are ordered by the median methylation level of tumor samples. **(c)** Boxplots displaying the transcriptional response to the demethylating agent 5-Aza-CdR in three HCC cell lines. The y-axis represents the log_2_ fold change of expression in treated cells versus DPBS controls. *P*-values were determined using two-sided Wilcoxon tests. **(d)** Stacked bar charts showing the chromosomal distribution of genes in the initial candidate pool (*n* = 1,294) and the final identified CTG set (*n* = 407). The *P*-value was determined by Fisher’s exact test comparing retention rates between autosomal and X-linked candidates (excluding Y-linked genes). **(e)** Scatter plot illustrating the relationship between median methylation level (MML; x-axis) and hypomethylation specificity index (HSI; y-axis) across all identified CTGs. The value for each gene was calculated as its median level across all analyzed TCGA cancer types. Spearman’s correlation and *P*-value are shown. Density plots display the distribution of MML (top) and HSI (right) for individual CTGs within corresponding CTG subgroups. Boxplots represent the median (center line), upper and lower quartiles (box limits), and 1.5 × IQR (whiskers). Significance: \**P* ≤ 0.05, \*\**P* ≤ 0.01, \*\*\**P* ≤ 0.001, \*\*\*\**P* ≤ 0.0001; ns, not significant. The definitions and color coding for CTG subgroups apply to panels **(a)**, **(c)** and **(e)**.

During CTG screening, X-linked genes were significantly underrepresented in the final CTG set (6.1%) relative to the initial candidate pool (14.5%), as confirmed by Fisher’s exact test (OR = 0.28; **Fig. 2d**). Intriguingly, 85 X-linked CTGs curated in the CTdatabase were absent from the final set, and this finding prompted a detailed examination of their expression profiles in malignant cells (**Supplementary Fig. 3c**). Among them, 13 showed undetectable expression in the Discovery Cohort; however, the remainder (exemplified by *CTAG1B)* were highly expressed in only a minority of tumor samples, with minimal or absent expression in the rest. These genes were therefore excluded, as our pipeline retained only genes recurrently activated in ≥50% of samples.

Tumor-associated DNA hypomethylation has been characterized as heterogeneous and stochastic rather than a uniform genome-wide loss^26–28^. Given the tight methylation dependency of X-linked CTGs, we hypothesized that their sample-specific activation reflects a similar pattern of DNA hypomethylation. To test this, we quantified two metrics for each CTG within each cancer type using TCGA methylation data: the median methylation level (MML) and the hypomethylation specificity index (HSI), which captures sample-specific promoter demethylation (see **Methods**). We then calculated the median values of these two metrics across cancer types and generated a scatter plot (**Fig. 2e**). The majority of X-linked CTGs mapped to a high-MML, high-HSI region, suggesting that their promoters are generally hypermethylated and that hypomethylation in a small subset of tumors underlies their sparse expression patterns. In contrast, autosomal CTGs were primarily distributed in the low-MML, low-HSI region, consistent with limited methylation-based regulation.

Taken together, these results indicate that X-linked CTGs are tightly regulated by DNA methylation and tend to show sample-specific expression due to stochastic demethylation of promoters.

### Chromatin regulators coordinate autosomal CTG activation

While the preceding analysis established DNA methylation as a dominant regulator of X-linked CTGs, the regulatory mechanisms driving activation of the broader CTG repertoire, particularly autosomal CTGs, remain to be delineated. To investigate this, we performed gene-gene co-expression analysis in malignant cells and found that the 407 CTGs exhibited significantly higher correlation with one another than non-CTG genes (**Fig. 3a**), indicating coordinated transcriptional regulation.

**Fig. 3.**
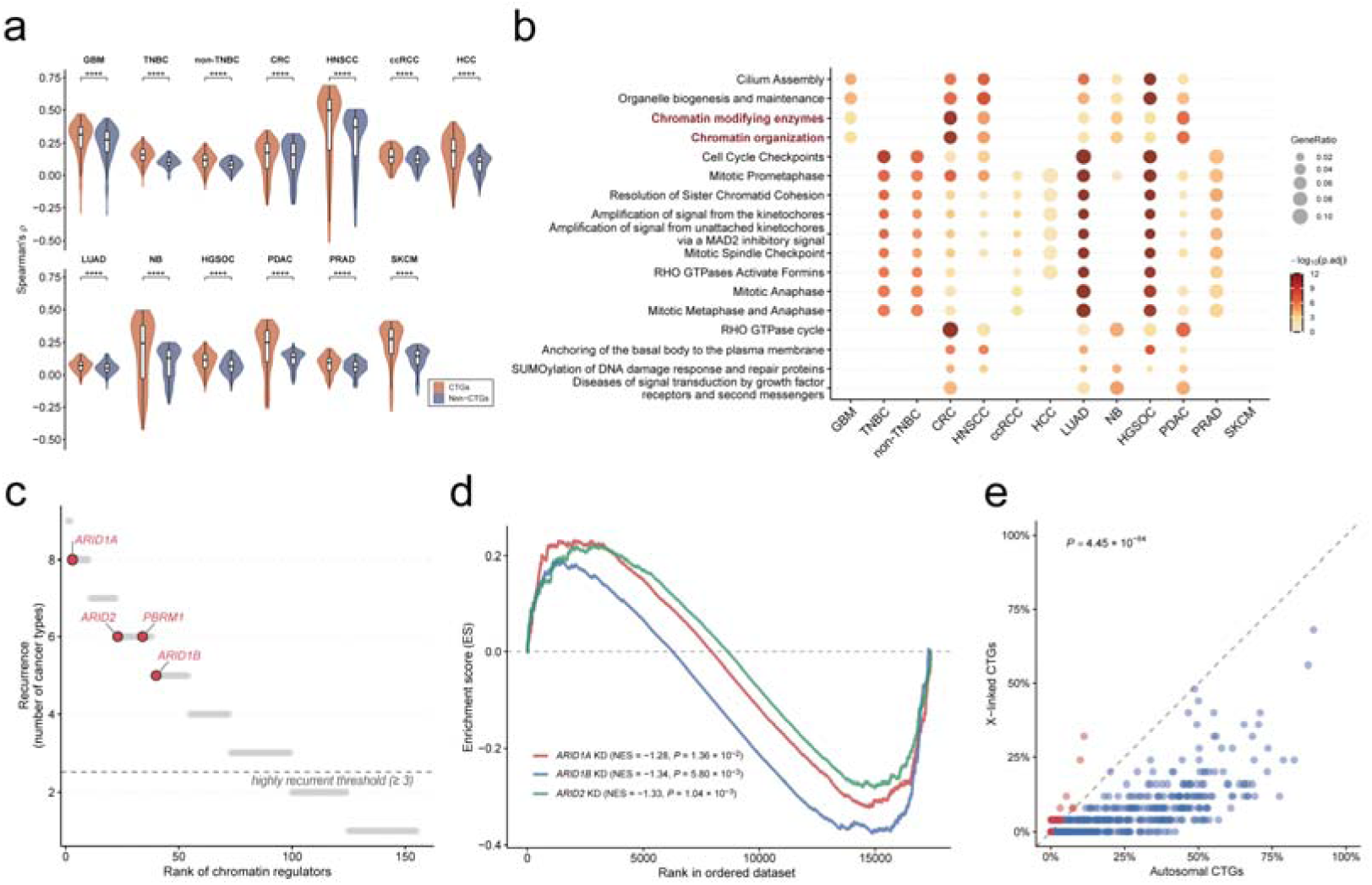
Chromatin regulators coordinate autosomal CTG activation. (a) Boxplots comparing the median Spearman correlation of individual CTGs with other CTGs versus with non-CTGs in the Discovery Cohort. *P*-values were determined using two-sided paired Wilcoxon signed-rank tests. (b) Reactome pathway enrichment analysis for the top 2,000 non-CTG genes exhibiting the highest frequency of significant positive co-expression with CTGs. The analysis was performed per cancer type using the Discovery Cohort. (c) Frequency-ranked candidate CTG regulators within the Reactome “Chromatin modifying enzymes” pathway across cancer types in the Discovery Cohort. (d) GSEA plots evaluating the global expression changes of the CTG signature in the transcriptomes of *ARID1A*-, *ARID1B*-, and *ARID2*-knockdown (KD) versus non-targeting control HepG2 cells. (e) Scatterplot comparing the proportion of regulator-occupied promoters between autosomal and X-linked CTGs. Each point represents an individual ChIP-seq dataset (*n* = 667). The grey dashed line indicates the line of *y* = *x*. The *P*-value was determined using a two-sided paired Wilcoxon signed-rank test. Boxplots represent the median (center line), upper and lower quartiles (box limits), and 1.5 × IQR (whiskers). Significance: \**P* ≤ 0.05, \*\**P* ≤ 0.01, \*\*\**P* ≤ 0.001, \*\*\*\**P* ≤ 0.0001; ns, not significant.

Motivated by this observation, for each cancer type in the Discovery Cohort, we ranked non-CTG genes based on the number of positively co-expressed CTGs and performed pathway enrichment analysis on the top 2,000 genes (**Fig. 3b**). Cell cycle-related pathways were universally enriched, suggesting a tight coupling between CTGs and cell-cycle programs. Notably, pathways associated with epigenetic regulation (including “chromatin modifying enzymes” and “chromatin organization”) were also significantly enriched in over half of the cancer types. Since the cell cycle itself is tightly regulated by epigenetic mechanisms^29,30^, we investigated whether this enrichment merely reflected a bystander effect. We therefore partitioned CTGs into cell cycle-associated and non-cell cycle-associated groups (see **Methods**) and repeated the analysis. Whereas the former showed the expected enrichment for mitotic pathways, the latter still retained significant enrichment for epigenetic pathways (**Supplementary Fig. 4a–b**).

To further identify specific regulators of CTG activation, we extracted genes annotated in the “chromatin modifying enzymes” pathway from the top-ranked gene list and prioritized them by recurrence across cancer types (**Fig. 3c**). In total, 99 genes were recurrently detected in at least three cancer types and retained as potential regulators, including genes encoding multiple core subunits of the SWI/SNF chromatin remodeling complex (e.g., *ARID1A*, *ARID1B*, *ARID2*, and *PBRM1*)^31^. Given that SWI/SNF-mediated nucleosome mobilization can increase chromatin accessibility and facilitate epigenetic modification^32,33^, we hypothesized that this complex may contribute to autosomal CTG activation. To functionally corroborate this, we analyzed a public bulk RNA-seq dataset from HepG2 hepatoblastoma cells following siRNA-mediated depletion of three biochemically distinct ARID subunits (*ARID1A*, *ARID1B*, and *ARID2*)^34^. Gene Set Enrichment Analysis (GSEA)^35^ revealed that depletion of any of these three subunits led to a significant downregulation of the CTG program (**Fig. 3d**), supporting a dependence of coordinated CTG activation on SWI/SNF complex activity.

In addition to gene expression data, we systematically analyzed chromatin immunoprecipitation followed by sequencing (ChIP-seq) data for these potential regulators from CistromeDB^36^ and calculated the proportion of CTG promoters directly occupied by them. Notably, the fraction of bound autosomal CTGs was significantly higher than that of X-linked CTGs (**Fig. 3e**). Collectively, these results indicate that autosomal CTGs are governed by an epigenetic mechanism distinct from that of X-linked CTGs and rely more heavily on the coordinated activity of specific chromatin regulators.

### Elevated CTG burden marks malignant cells in single-cell transcriptomes

Accurate annotation of malignant cells is a foundational step in the analysis of tumor scRNA-seq data. Building on our observation that CTG expression effectively distinguishes bulk tumor from normal samples, we investigated whether CTG activation at single-cell resolution also constitutes a robust hallmark of malignancy. Given the sparsity of single-cell transcriptomic data, aggregated CTG expression may be disproportionately influenced by a limited subset of highly expressed genes. We therefore introduced CTG burden, defined as the number of expressed CTGs (denoted as *N*_CTG_), to quantify CTG activation across single cells. Since CTG burden derived from the full 407 CTGs was highly concordant with that based on CTGs identified in corresponding individual cancers (**Supplementary Fig. 5a**), we employed the complete CTG set for all subsequent analyses to ensure consistency.

Across all 13 cancer types in the Discovery Cohort, malignant cells consistently showed significantly higher CTG burden than non-malignant cells (**Supplementary Fig. 5b**). Notably, this elevation persisted when non-malignant cells were further stratified into five major categories (immune, stromal, epithelial, glial, and melanocytic cells; **Fig. 4a** and **Supplementary Fig. 5c**). To confirm that such elevation reflected malignant transformation rather than lineage differences, we specifically compared CTG burden between malignant cells and cell-type-matched normal references (e.g., epithelial cells for carcinomas, glial cells for GBM). In the majority of samples, malignant cells still exhibited significantly higher CTG burden than their corresponding normal reference cells (**Supplementary Fig. 5d**), indicating that elevated CTG burden is a robust feature of malignancy.

**Fig. 4.**
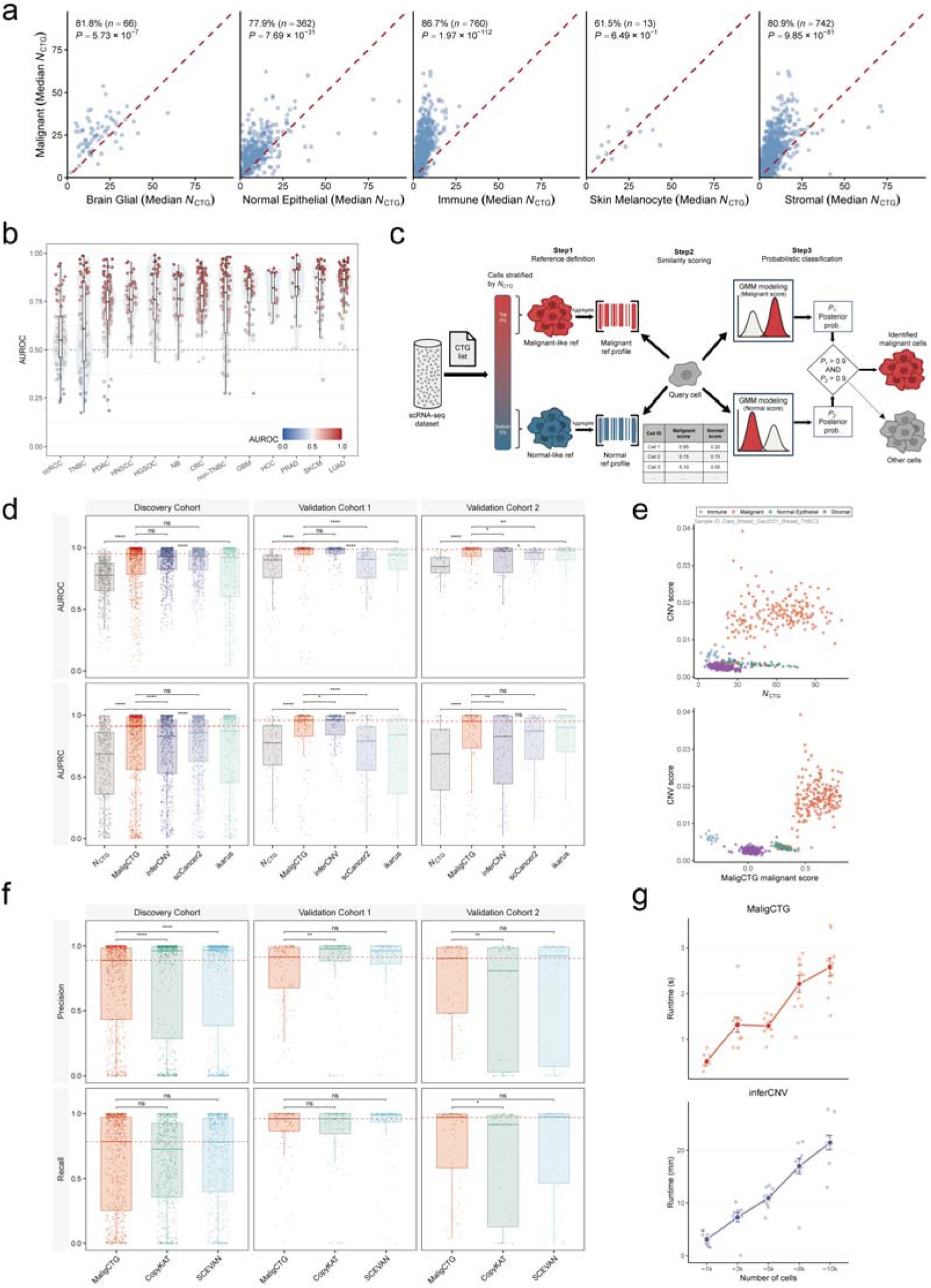
Elevated CTG burden marks malignant cells in single-cell transcriptomes. **(a)** Pairwise comparisons of the median *N*_CTG_ between malignant cells (y-axis) and five distinct non-malignant cell populations (x-axis) within the same samples. Each dot represents an individual tumor sample from the Discovery Cohort. The red dashed line represents the line of *y* = *x*. The percentages indicate the proportion of samples wherein malignant cells harbor a higher median *N*_CTG_ than non-malignant cells. *P*-values were determined using two-sided paired Wilcoxon signed-rank tests. **(b)** Violin plots of AUROC values quantifying the distinction in *N*_CTG_ levels between malignant and non-malignant cells across samples in the Discovery Cohort. **(c)** Schematic overview of the MaligCTG pipeline. The workflow involves defining reference profiles based on extreme *N*_CTG_ values, computing transcriptomic similarity scores, and assigning final binary labels using a Gaussian mixture model (GMM). **(d)** Boxplots comparing the continuous scoring accuracy of MaligCTG against *N*_CTG_, inferCNV, scCancer2, and ikarus across three cohorts. Performance was evaluated using AUROC and the area under the precision-recall curve (AUPRC). This analysis includes malignant cells and all non-malignant cells. **(e)** Scatterplots demonstrating the improved distinction of malignant cells from non-malignant cells using MaligCTG (bottom) compared with *N*_CTG_ (top). The x-axis represents the respective scores, plotted against inferCNV-derived CNV scores (y-axis) for individual cells in a representative sample. **(f)** Boxplots comparing the binary classification performance of MaligCTG against CopyKAT and SCEVAN. Evaluation metrics include precision and recall. This analysis includes malignant cells and all non-malignant cells. **(g)** Line plots illustrating the runtime of MaligCTG (top) and inferCNV (bottom) across samples containing varying numbers of cells. For each cell-number bin, 10 samples were randomly selected. Results are shown as mean ± standard error (SE). *P*-values were determined using two-sided Wilcoxon tests for comparisons in panels **(d)** and **(f)**. Boxplots represent the median (center line), upper and lower quartiles (box limits), and 1.5 × IQR (whiskers).

Prompted by this observation, we next evaluated the utility of CTG burden for malignant cell annotation and achieved a median AUROC ≥ 0.75 in 11 of 13 cancer types in the Discovery Cohort (**Fig. 4b**). Interestingly, the median AUROC was positively correlated with the number of identified CTGs across cancer types (**Supplementary Fig. 5e**), and ccRCC exhibited the lowest performance (median AUROC = 0.55), consistent with its CTG-poor phenotype. In addition to CTG burden, we also utilized aggregated CTG expression as a feature for malignant cell annotation and found its performance was inferior to that of CTG burden, suggesting CTG burden is a more informative summary metric for distinguishing malignant cells (**Supplementary Fig. 5f**).

Although CTG burden alone already provided excellent performance, we further developed a pipeline termed MaligCTG to optimize malignancy annotation (**Fig. 4c**). Briefly, given a cancer single-cell dataset, MaligCTG first defines putative transcriptomic profiles of malignant and normal cells based on cells with extreme CTG burden (top and bottom 5%, respectively). Transcriptomic similarity scores are then computed for all cells relative to these reference profiles, yielding continuous malignant and normal scores. As expected, the malignant score derived from MaligCTG outperformed CTG burden in the identification of malignant cells (**Fig. 4d**), and **Fig. 4e** illustrates this enhancement in a representative TNBC sample. MaligCTG also supports binary classification by using a two-component Gaussian mixture model (GMM) to assign each cell as malignant or non-malignant.

We next benchmarked MaligCTG against five established malignancy annotation tools using samples from the Discovery Cohort. For methods that output continuous values (including inferCNV^37^, scCancer2^38^, and ikarus^39^), MaligCTG demonstrated comparable performance to inferCNV and slightly outperformed scCancer2 and ikarus (**Fig. 4d**). For methods that output binary labels (including CopyKAT^40^ and SCEVAN^41^), MaligCTG also achieved comparable performance (**Fig. 4f**). To assess generalizability, we further benchmarked the method in two additional cohorts: Validation Cohort 1, consisting of 172 tumor samples sequenced on Smart-seq2 and spanning nine cancer types represented in the Discovery Cohort; and Validation Cohort 2, consisting of 81 tumor samples from 12 independent cancer types absent from the Discovery Cohort (**Fig. 1a and Supplementary Fig. 1a–c**). As expected, no significant performance drop was observed for MaligCTG, underscoring its robustness (**Fig. 4d** and **4f**). Importantly, this robust performance persisted even when the evaluation was confined to malignant cells and their corresponding normal reference populations (**Supplementary Fig. 6a–b**).

MaligCTG not only achieved annotation accuracy comparable to that of widely used methods such as inferCNV, but also offered substantial computational acceleration (**Fig. 4g**). For instance, for datasets with ∼5,000 cells, inferCNV required approximately 11 minutes to finish, whereas MaligCTG completed the task in only 1.3 seconds. Such efficiency makes MaligCTG particularly suitable for large-scale malignancy annotation in atlas-level studies.

Collectively, our results suggest that CTG burden constitutes a robust hallmark of malignancy in single-cell transcriptomic data, and we present MaligCTG as a rapid and accurate pan-cancer framework for malignant-cell annotation based on the identified CTGs.

### Distinct spatial organization of CTG activation across cancer types

While our single-cell analysis elucidated the transcriptional characteristics of CTGs, their spatial organization within tumors remained unresolved. To explore this question, we curated 245 spatial transcriptomic (ST) slides spanning 15 cancer types (**Fig. 1a**). We utilized CTG burden to quantify CTG activation for each spot and observed that it exhibited a substantial positive correlation with tumor purity (Spearman’s ρ > 0.5) in 10 of 15 cancer types (**Fig. 5a–b**), suggesting that CTG burden may serve as a surrogate for purity. Consistent with this association, CTG burden was significantly higher in malignant spots than in other cell types in two well-annotated ST datasets^42,43^ (**Supplementary Fig. 7a–b**).

**Fig. 5.**
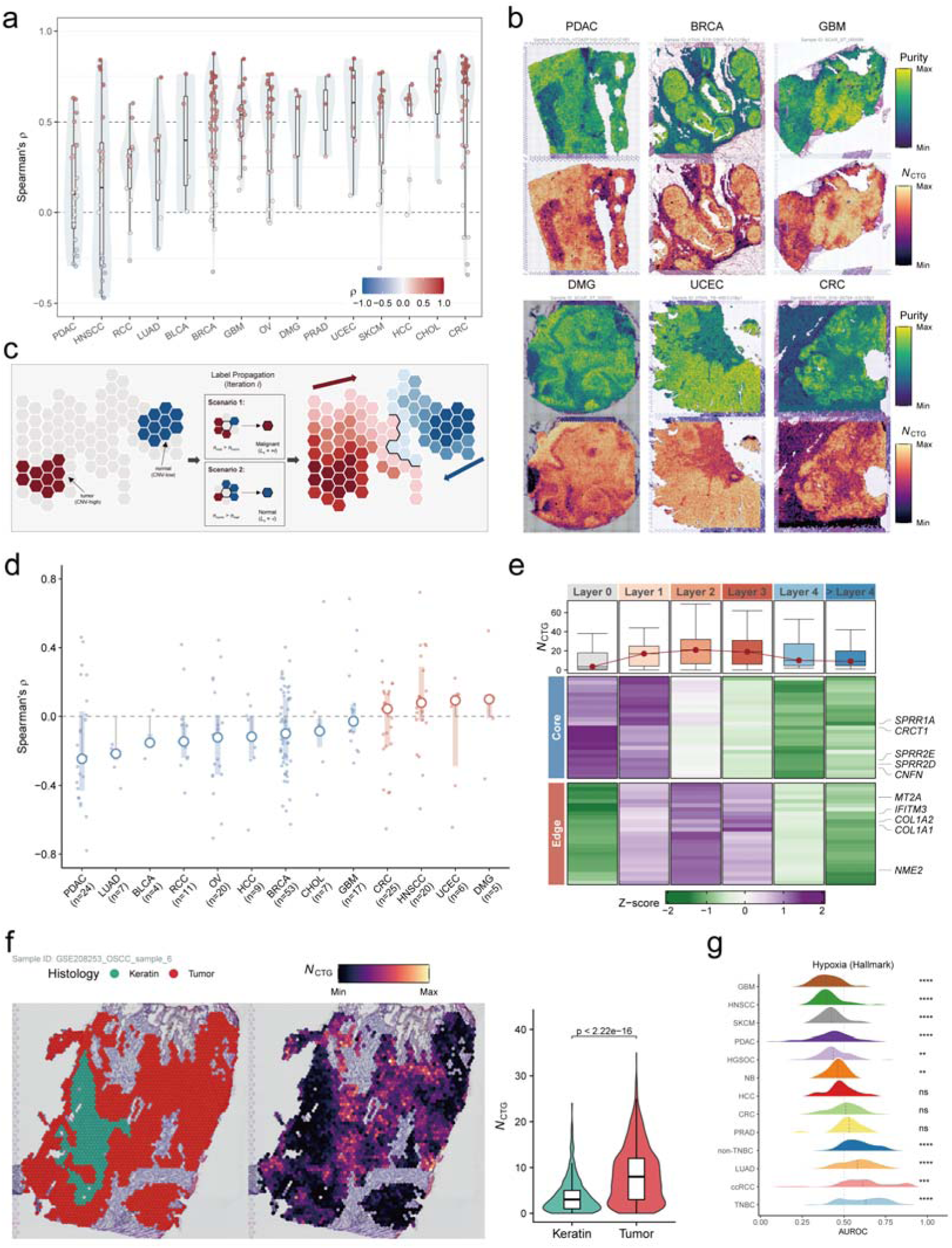
Distinct spatial organization of CTG activation across cancer types. **(a)** Violin plots showing the Spearman correlation coefficients between *N*_CTG_ and tumor purity across spatial transcriptomics (ST) samples from diverse cancer types. Each point represents an individual sample. **(b)** Spatial visualization of estimated tumor purity and *N*_CTG_ levels in six representative ST slices. **(c)** Schematic illustration of the label propagation algorithm used to model the tumor–stroma transition. **(d)** Lollipop plot summarizing the Spearman correlation between purity-adjusted *N*_CTG_ levels and tumor layer indices across cancer types. Large dots represent the median Spearman correlation, with error bars denoting the IQR. Small jittered points represent individual samples. Colors indicate whether the median correlation was positive or negative. **(e)** Heatmap illustrating the expression of edge-associated and core-associated gene signatures along the propagated tumor layers for a representative HNSCC sample. The upper box plots depict the corresponding *N*_CTG_ values along the layers. Representative genes are labeled on the right. **(f)** Spatial visualization of pathologist-annotated keratin and tumor regions (left) and *N*_CTG_ values in these regions (middle). The accompanying violin plot (right) quantifies the difference in *N*_CTG_ levels between tumor and keratin regions. The *P*-value was computed using a two-sided Wilcoxon rank-sum test. **(g)** Ridge plot showing the distribution of AUROC values for hypoxia pathway activity across cancer types. An AUROC value > 0.5 indicates elevated hypoxia activity in CTG-high malignant cells within a sample, whereas a value < 0.5 indicates higher activity in CTG-low cells. The gray dashed line indicates the baseline value of 0.5. Vertical black dashed lines indicate the median for each cancer type. *P*-values were computed using a two-sided one-sample Wilcoxon signed-rank test against a theoretical median of 0.5. Significance: \**P* ≤ 0.05, \*\**P* ≤ 0.01, \*\*\**P* ≤ 0.001, \*\*\*\**P* ≤ 0.0001; ns, not significant. Abbreviations: OV, ovarian cancer; DMG, diffuse midline glioma; BLCA, bladder urothelial carcinoma; CHOL, cholangiocarcinoma; UCEC, uterine corpus endometrial carcinoma.

We next developed an iterative neighbor-voting algorithm to reconstruct the tumor–stroma boundary (**Fig. 5c**). Briefly, we defined high-confidence malignant spots as tumor cores and high-confidence normal spots as stromal regions based on copy-number variation (CNV) profiles. These spots were assigned initial “tumor” and “stroma” labels, respectively, which were then iteratively propagated to adjacent unlabeled spots until all spots were labeled. For each “tumor” spot, we recorded the iteration at which the label was assigned and defined this value as the layer index; a higher layer index therefore indicates increasing distance from the tumor core toward the surrounding stroma.

Leveraging this layered representation, we examined the relationship between CTG burden and propagated tumor layers. To control for the influence of tumor purity, we analyzed the association between purity-adjusted CTG burden and layer index (**Fig. 5d**). In nine cancer types, the median correlation value was negative, suggesting that CTG activation may generally be higher in the tumor core. By contrast, four cancer types showed a positive median correlation. Among them, HNSCC was of particular interest because 65% of the ST slides exhibited a positive correlation, indicating that CTG activation may be enriched in the tumor boundary region. This spatial pattern was illustrated by a representative HNSCC sample, in which CTG burden across tumor layers closely tracked the expression of previously defined edge-associated genes and showed the opposite trend relative to core-associated genes^42^ (**Fig. 5e**). Analysis of four HNSCC slides containing pathologist-annotated keratin regions further showed that CTG activation tended to localize outside keratinized tumor regions, which are typically situated in the core of cancer nests, although the magnitude of this difference varied across samples (**Fig. 5f and Supplementary Fig. 8a–c**).

Hypoxia is widely recognized as a feature of tumor cores^44^, and we therefore compared hypoxia activity between malignant cells with high and low CTG burden using single-cell data from the Discovery Cohort (see **Methods**). In most HNSCC samples, malignant cells with lower CTG burden showed higher hypoxia scores; by contrast, in cancers such as TNBC and ccRCC, hypoxia levels were significantly higher in cells with high CTG burden (**Fig. 5g)**. Taken together, these results revealed the spatial heterogeneity of CTG activation. Although CTG activation was generally enriched in tumor cores across most cancer types, HNSCC showed preferential activation at the tumor–stroma boundary.

### Prioritization of CTGs as candidate targets for TCR-T therapy

T-cell receptor-engineered T-cell (TCR-T) therapy is a promising therapeutic modality that requires tumor-restricted antigens to minimize off-tumor toxicity^45^. The restricted expression of CTGs in testicular germ cells renders them inherently attractive TCR-T targets, a rationale underscored by Tecelra^46^ (afamitresgene autoleucel), the first FDA-approved TCR-T therapy for synovial sarcoma. However, existing CTG-directed TCR-T trials have largely been confined to canonical cancer-testis antigens. Building on our high-confidence CTG catalog, we therefore sought to expand the repertoire of candidate TCR-T targets. To this end, candidate targets were required to satisfy three criteria: (i) no association with the cell cycle, to reduce the risk of on-target, off-tumor toxicity in proliferative normal tissues; (ii) negligible expression in non-testicular normal tissues; and (iii) robust upregulation in malignant cells. This strategy yielded 10 candidates from the 407 CTGs, which were reduced to nine after exclusion of one long non-coding RNA (**Fig. 6a**). Among them, *ROPN1* was noteworthy because it had previously been reported as a TCR-T target showing substantial efficacy in TNBC mouse models^47^. Its recovery in our prioritization analysis provides independent support for the robustness of this pipeline.

**Fig. 6.**
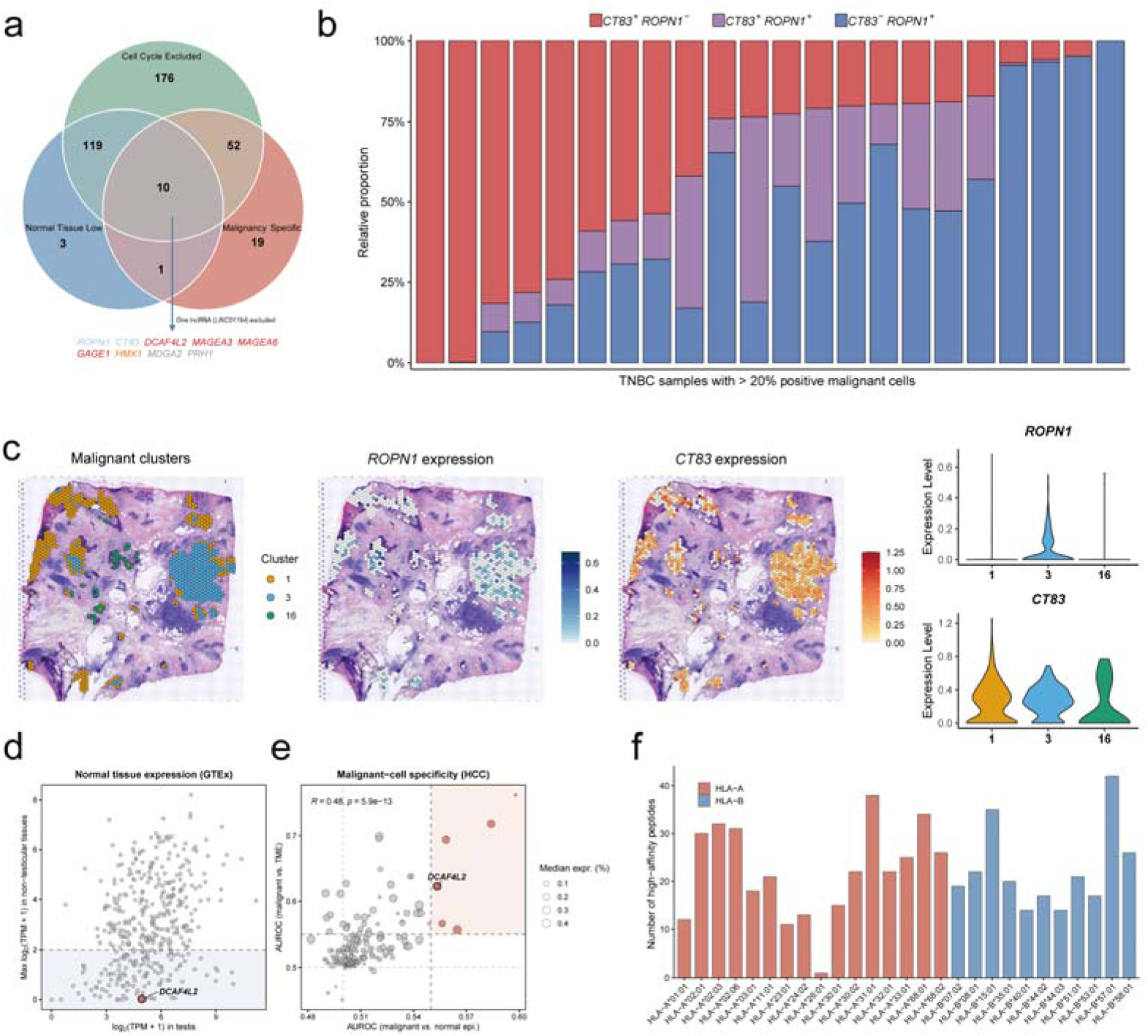
Prioritization of CTGs as candidate targets for TCR-T therapy. **(a)** Venn diagram illustrating the overlap of genes meeting three prioritization criteria: cell cycle exclusion, low normal tissue expression, and malignancy-specific expression. The nine final candidates are listed and color-coded according to the cancer type in which they were prioritized. **(b)** Stacked bar plot depicting the relative proportions of *CT83*^+^*ROPN1*^−^, *CT83*^+^*ROPN1*^+^, and *CT83*^−^*ROPN1*^+^ subpopulations among malignant cells for each TNBC sample from the Discovery Cohort in which >20% of malignant cells expressed at least one of these genes. **(c)** Spatial transcriptomic visualization of malignant clusters (left), *ROPN1* expression (middle), and *CT83* expression (right) in a representative breast cancer tissue section. The accompanying violin plots depict the expression levels of *ROPN1* and *CT83* across identified malignant clusters. **(d)** Scatter plot illustrating the expression of CTGs in normal tissues based on the GTEx dataset; the y-axis displays the maximum expression across all non-testicular normal tissues, and the x-axis displays the expression in testis. The light blue shaded region indicates the CTGs with maximum non-testicular expression ≤ 3 TPM. **(e)** Scatter plot demonstrating the malignant-cell specificity of CTGs in HCC at single-cell resolution. Axes represent the AUROC values for distinguishing malignant cells from normal epithelial cells (x-axis) and the TME cells (y-axis). The light red shaded region outlines CTGs with AUROC values ≥ 0.55 in both axes. Bubble size reflects the median expression percentage of each CTG in malignant cells. Spearman correlation coefficient and *P*-value are shown. **(f)** Bar plot enumerating the predicted high-affinity DCAF4L2-derived peptides across MHC class I alleles. Significance: \**P* ≤ 0.05, \*\**P* ≤ 0.01, \*\*\**P* ≤ 0.001, \*\*\*\**P* ≤ 0.0001; ns, not significant.

Among the remaining eight genes, two gained our particular interest. The first was *CT83* (also known as *KK-LC-1*), which we prioritized as a candidate TCR-T target for TNBC. Given the promising results of *ROPN1*-targeted TCR-T therapy, we examined whether *CT83* represents an independent target distinct from *ROPN1*. For each TNBC sample in the Discovery Cohort in which more than 20% of malignant cells expressed one or both genes, we calculated the proportion of *CT83*^+^*ROPN1*^−^, *CT83*^+^*ROPN1*^+^ and *CT83*^−^*ROPN1*^+^ subpopulations (**Fig. 6b**). Notably, in 8 of 22 samples, the percentage of *CT83*^+^*ROPN1*^−^ malignant cells exceeded 50%. In addition, we examined spatial transcriptomics data from breast cancer. In a representative sample, spot-level clustering identified three malignant clusters: clusters 1 and 16 preferentially expressed *CT83* alone, whereas cluster 3 co-expressed *CT83* and *ROPN1* (**Fig. 6c and Supplementary Fig. 9a–b**). Importantly, these clusters occupied spatially distinct regions within the tumor. Together, these data suggest that *CT83* and *ROPN1* mark partially non-overlapping TNBC malignant cell populations, and that their combinatorial targeting might offer a rational strategy to address intratumoral heterogeneity and potentially enhance the efficacy of TCR-T therapy.

The second candidate was *DCAF4L2*, an autosomal CTG prioritized as a candidate TCR-T target for liver cancer. According to GTEx, *DCAF4L2* showed robust expression in the testis but extremely low expression across somatic tissues, suggesting a favorable safety profile (**Fig. 6d**). Differential expression analysis revealed marked *DCAF4L2* reactivation in HCC malignant cells; it was expressed in over 20% of malignant cells while remaining largely absent from normal epithelial cells and tumor microenvironment (TME) cells (**Fig. 6e**). To assess its antigenic potential, we utilized MHCflurry^48^ to predict HLA class I-restricted binding peptides and identified 258 high-affinity candidates derived from *DCAF4L2* across 27 prevalent MHC class I alleles (e.g., HLA-A*02:01) (**Fig. 6f**). We also queried the Immune Epitope Database (IEDB)^49^ and retrieved five experimental epitope records. Notably, one of these epitopes (LSHDSAVTSL), which MHCflurry predicted to bind strongly to HLA-B*57:01, had also been detected by mass spectrometry elution in malignant cells from HCC patients.

Collectively, our results highlight the underexplored therapeutic potential of CTGs in cancer treatment. Although neither *CT83* nor *DCAF4L2* belongs to the most extensively studied CTGs such as *MAGEA4* and *NY-ESO-1*, our analysis suggests that they represent plausible TCR-T candidate targets and warrant further exploration.

## Discussion

CTGs have long attracted interest as immunotherapy targets because they are normally restricted to the testis and are aberrantly reactivated in tumors^1–4^. A pan-cancer CTG catalog is therefore essential for translating this class of antigens into clinically actionable strategies. However, when bulk RNA-seq data are used to identify CTGs, it is often difficult to determine whether the detected expression of a CTG candidate reflects reactivation in malignant cells or contamination from normal cells^50^. By constructing our catalog from single-cell transcriptomic data, we ensured that the identified CTGs exhibit cancer-cell-intrinsic activation. Although the 10x Genomics platform may underdetect low-abundance transcripts and thereby miss a subset of CTGs, independent validation using GTEx and TCGA data supported the overall quality of the catalog.

One fundamental question in CTG biology concerns the mechanism governing their cancer-specific reactivation. While promoter DNA demethylation is canonically viewed as the primary “on-switch” for CTG activation^22–24^, our multi-omics analysis revealed a heterogeneous regulatory landscape. We found that methylation dependency is largely confined to X-linked CTGs, which consequently exhibited stochastic and sample-restricted expression patterns. In contrast, autosomal CTGs showed broader activation across tumor samples and appeared to be governed by extensive networks of chromatin-modifying enzymes, including the SWI/SNF remodeling complex. Consistently, our ChIP-seq analysis showed that promoters of autosomal CTGs were more permissive to direct occupancy by these chromatin regulators than were X-linked CTG promoters. The mechanisms underlying this distinct regulatory dependency remain unclear and warrant further investigation.

Despite the sparsity of single-cell transcriptomic data and the generally low expression of individual CTGs, we found that elevated CTG burden is a robust pan-cancer hallmark of malignancy. Building on this observation, we developed MaligCTG, a computational pipeline for malignant-cell annotation. Compared with the widely used inferCNV, which infers malignancy from copy number variation, MaligCTG offers two practical advantages. First, it is computationally efficient because it avoids genome-wide sliding-window smoothing of expression profiles, enabling annotation within seconds rather than minutes for the same task in our benchmarking. Second, it does not require prior selection of diploid reference cells, a step that is indispensable for inferCNV and often nontrivial in practice. Together, these features make MaligCTG particularly useful for integrative analyses of large-scale cancer single-cell transcriptomic datasets in which computational cost is a bottleneck.

Our findings also have immediate implications for the design of TCR-T therapies. First, current strategies largely focus on X-linked CTGs such as *MAGEA4* and *NY-ESO-1*^3,4^; however, our results demonstrate that the expression of these genes is often restricted to a small fraction of tumor samples, which may explain the variable clinical efficacy observed in prior trials^51,52^. Given the methylation dependency of X-linked CTGs, DNMT inhibition may provide a rationale for increasing their expression and thereby improving the feasibility of CTG-directed immunotherapy. Second, to overcome the limitation of sparse antigen expression, our study selected for CTGs that are broadly activated across tumor samples. In this context, our prioritization framework yields a new panel of candidate TCR-T target genes, and the recovery of *ROPN1* provides internal support for its validity. Notably, *CT83* emerged as a complementary TCR-T target in TNBC, where its expression only partially overlapped with that of *ROPN1*, suggesting that combinatorial targeting may help address intratumoral heterogeneity. Third, CTG-directed TCR-T may be particularly relevant in HCC, a malignancy in which the current TCR-T literature remains dominated by hepatitis B virus (HBV)- and alpha-fetoprotein (AFP)-directed approaches^53^. Our analysis indicated that X-linked CTGs are highly activated in this cancer type, rendering HCC particularly amenable to CTG-directed TCR-T therapy. Beyond these canonical X-linked CTGs, we identified the autosomal CTG *DCAF4L2* as a candidate TCR-T target for HCC, which merits further exploration.

Two limitations should be considered in future studies. First, because our analyses were primarily based on primary and metastatic tumor specimens, CTGs selectively activated in circulating tumor cells or premalignant lesions may have been overlooked. Such genes may be informative for early detection and should be considered in future catalog refinement. Second, most cancer types included in this study were adult cancers, and it remains unclear to what extent these findings generalize to pediatric cancers.

In conclusion, our comprehensive pan-cancer analysis deepens our understanding of CTG biology and supports the development of novel CTG-targeted therapeutic strategies. As additional single-cell and multi-omic datasets become available across diverse cohorts and platforms, iterative catalog refinement will further support the translational development of CTG-based therapies.

## Methods

### Datasets collection

A total of 90 well-annotated cancer single-cell and single-nucleus RNA sequencing (scRNA-seq and snRNA-seq) datasets were collected primarily from the Single Cell Portal (https://singlecell.broadinstitute.org/single_cell), 3CA^13^, HTAN^14^, and TISCH2^15^. For each dataset *d*, the following criteria were applied: (1) if *d* encompassed samples from distinct cancer subtypes or sequencing platforms, then it was stratified into separate datasets; (2) only tumor samples containing ≥10 malignant cells and adjacent normal samples were retained; and (3) specimens obtained from the same patient at multiple timepoints (e.g., pre- and post-treatment) or anatomical sites (e.g., tumor core and edge) were treated as independent samples when sufficient metadata were available. The resulting datasets were organized into one discovery cohort and two independent validation cohorts.

Spatial Transcriptomics (ST) datasets comprising 245 tumor sections across 15 cancer types were acquired from GEO, SCAR^16^, HTAN, Mendeley Data, and CROST^17^.

Bulk RNA-seq data from TCGA, TARGET, and GTEx projects were downloaded from UCSC Xena (https://xenabrowser.net/datapages/). DNA methylation data for TCGA samples generated using the Illumina HumanMethylation450 BeadChip platform were also obtained from UCSC Xena. RNA-seq data from HCC cell lines (HepG2, SNU398, and SNU475) treated with the demethylating agent 5-Aza-CdR were retrieved from GEO (GSE202560). RNA-seq data from gene knockdown experiments were downloaded from GEO (GSE69567). ChIP-seq data for these recurrent regulators were downloaded from CistromeDB^36^.

### Assembly of CTG candidate genes

A total of 276 genes were retrieved from the CTdatabase (http://www.cta.lncc.br/). After removal of genes lacking approved HGNC symbols and corresponding Ensembl IDs, 242 genes were retained. In parallel, testis-enriched protein-coding genes were extracted from the Human Protein Atlas (HPA; https://www.proteinatlas.org/). Genes were retained if their expression level in testis or a tissue group (two to five tissues including testis) was at least fourfold higher than in any other tissue. A total of 1,212 genes with approved HGNC symbols met this criterion. The union of these two sources yielded 1,294 unique CTG candidates.

### Quantification of the aggregated expression level of a CTG set

For bulk RNA-seq samples, the aggregated expression level of the CTG set (*E*_CTG_) was quantified using single-sample Gene Set Enrichment Analysis (ssGSEA)^54^. For single-cell data, *E*_CTG_ was calculated using the Seurat *AddModuleScore* function^55^.

### Cell type harmonization and definition of normal reference cells

Original cell type annotations provided within each dataset were harmonized and reclassified into eight major lineages: normal epithelial, stromal, immune, brain glial, skin melanocyte, malignant, other (e.g., adipocytes and myocytes), and unknown (e.g., low-quality cells). Normal reference cells were defined as non-malignant cells of the same lineage origin as the corresponding tumor type (e.g., epithelial cells for carcinomas, glial cells for GBM). Reference cells were restricted to those derived from primary tumor or normal tissue samples; cells from metastatic lesions were excluded.

### Inference of copy number variations in sc/snRNA-seq data

Copy number variations (CNVs) in individual cells were inferred at the sample level using the infercnv R package (v1.17.0). Reference cells were defined as the stromal or immune population containing the largest number of cells; when only binary annotations were available (malignant or non-malignant), all non-malignant cells were designated as reference cells. Gene-level CNV estimates were derived by centering the residual expression values (subtracting 1) to set the diploid baseline to zero. A global CNV score was subsequently calculated for each cell as the mean of the squared gene-level CNV values.

### Calculation of hypomethylation specificity index (HSI)

To quantify sample-specific promoter hypomethylation, we defined the hypomethylation specificity index (HSI) based on the statistical skewness of the promoter hypomethylation distribution. For a given gene *g* in sample *i*, the promoter DNA methylation level (β*_i_)* was defined as the average β-value across all Illumina array probes mapping to its promoter regions, including TSS200, TSS1500, 5’UTR, and the first exon. The corresponding promoter hypomethylation level (*x_i_*) was subsequently derived as *x_i_*= 1 - β*_i_*. For a given cancer type *C* comprising *N* samples, the HSI of gene *g* was computed as follows:

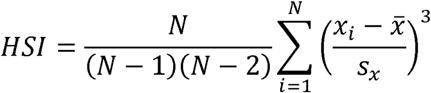

where x_□_ and *s_x_* denote the sample mean and sample standard deviation of the hypomethylation levels for gene *g* across the *N* samples within the cohort, respectively. A larger positive HSI value indicates a right-skewed distribution, reflecting sample-specific hypomethylation events confined to a minority of samples within a given cancer type.

### Gene-gene co-expression analysis in malignant cells

For each tumor sample in the Discovery Cohort, expression values from all malignant cells were aggregated by applying Seurat’s *AggregateExpression* function to produce a pseudoLbulk profile per sample. Given a cancer type *C* and two genes, Spearman-rank correlation was computed across pseudo-bulk profiles of samples of *C*.

To investigate pathways associated with CTG activation, we first extracted CTG and non-CTG pair which exhibited strong positive co-expression (ρ > 0.5 and adjusted *P*-value < 0.01). Non-CTGs were then ranked by the number of CTGs with which they exhibited strong positive correlation. The top-ranked genes per cancer type were subjected to pathway enrichment analysis against the Reactome database^56^ using the *compareCluster* function in clusterProfiler^57^ (v4.6.2).

### Definition of cell cycle-associated CTGs

To pinpoint CTGs linked to proliferation, we first quantified cell cycle activity in malignant cells using AUCell (v1.20.2)^58^ with G2/M and S phase gene sets^59^. Malignant cells were classified as “cycling” if their AUCell score exceeded 5 × MAD (median absolute deviations) for either phase. We then performed differential expression analysis between cycling and non-cycling cells using a Wilcoxon rank-sum test (Seurat *FindMarkers*, pseudocount = 10^-9^) within each dataset. For a given cancer type, CTGs were defined as cell cycle-associated if they were significantly upregulated (adjusted *P*-value < 0.01, average logL Fold Change > 1, and a difference in the fraction of expressing cells > 0.1) across ≥ 50% of datasets (or ≥ 2 datasets for cancer types with fewer than three datasets). All cell cycle-associated CTGs were pulled together considering their similarity between cancer types.

### The MaligCTG algorithm

For a given single-cell transcriptomic dataset (or sample), cells were stratified by CTG burden (*N*_CTG_), and the top and bottom 5% were designated as putative malignant and normal reference populations, respectively. Pseudo-bulk reference profiles were then generated for the two populations. Using the SingleR package (v2.0.0)^60^, transcriptomic similarity to the malignant and normal reference profiles was quantified for each cell, yielding malignant and normal scores, respectively.

Both score distributions were independently modeled using two-component Gaussian mixture models (GMMs). For the malignant-score GMM, the component with the higher mean was defined as the malignancy-associated state; conversely, for the normal-score GMM, the component with the lower mean was designated as the malignancy-associated state. Let *P_1_* and *P_2_* denote the posterior probabilities of a cell belonging to these respective malignancy-associated components. To ensure high specificity, cells were classified as malignant only when both probabilities exceeded 0.9 (*P*L > 0.9 and *P*L > 0.9).

### Benchmarking the performance of MaligCTG for malignant cell identification

The predictive performance of MaligCTG was benchmarked against five established methods for malignant cell identification. When comparing with methods whose outputs were continuous scores, the area under the receiver operating characteristic curve (AUROC) and the area under the precision-recall curve (AURPC) were calculated for each method. When comparing with methods whose outputs were binary labels, precision and recall were used as performance metrics. All benchmarking analyses were conducted at the sample level, and only samples containing at least 10 malignant cells and 10 non-malignant cells were included.

Comparative methods were implemented as follows: (1) for infercnv, CNV scores were derived as detailed in the “**Inference of copy number variations in sc/snRNA-seq data**” section; (2) for ikarus, malignancy probabilities were generated from raw counts using default parameters; (3) for scCancer2, the pre-trained XGBoost classifier was applied to scaled expression matrices comprising the top 5,000 highly variable genes, yielding probabilities of malignancy; and (4) for CopyKAT and SCEVAN, discrete binary malignant labels were independently inferred using the copykat (v1.1.0) and SCEVAN (v1.0.3) packages, respectively, with default parameters.

### Comparison of hypoxia activity between CTG-high and CTG-low malignant cells

For each sample, an adaptive threshold was defined as the 99th percentile of *N*_CTG_ observed in non-malignant tumor microenvironment cells. Malignant cells exceeding this threshold were classified as the CTG-high subpopulation, whereas the remainder were designated as CTG-low. Hypoxia activity scores for individual cells were quantified using the Seurat *AddModuleScore* function; the “Hypoxia” gene set from the MSigDB Hallmark collection^61^ was used in this calculation.

Within each sample, the hypoxia activity score was employed as a classification feature to discriminate CTG-high from CTG-low cells, and the AUROC was calculated to quantify the difference in hypoxia activity between the two subpopulations. An AUROC value higher than 0.5 indicates higher hypoxia activity in CTG-high malignant cells, and *vice versa*. For each cancer type, sample-level AUROC values were aggregated, and overall pathway alteration was summarized using the median AUROC. Statistical significance was assessed using a two-sided one-sample Wilcoxon signed-rank test against a theoretical median of 0.5 to evaluate the overall difference in hypoxia activity between the two subpopulations within a given cancer type.

### Spatial tumor layering based on inferred copy number variations

To characterize tumor spatial organization, raw ST count matrices were processed using the *SCTransform* pipeline, followed by dimensionality reduction and transcriptome-based Louvain clustering (resolution = 1.5) via the Seurat R package.

Spatially resolved copy number variations (CNVs) were inferred using the framework described by Xun et al^62^. To establish normal reference spots, transcriptome-based clusters were evaluated for canonical immune marker expression (e.g., *PTPRC*, *CD3D*, and *MS4A1*); only clusters showing significant immune enrichment (AUROC > 0.6 and adjusted *P* < 0.01 by Wilcoxon rank-sum test) were retained. The algorithm was then applied using random tree partitioning to stratify spots into eight CNV-defined clusters. Genome-wide CNV scores were calculated as the sum of absolute deviations from the diploid baseline across all genes. Among these, the two clusters with the highest median CNV scores were designated as CNV-high (malignant tumor cores), whereas the two with the lowest median CNV scores were designated as CNV-low (non-malignant stromal barriers).

Using these CNV-based seeds, we developed a constrained iterative label propagation algorithm to model the spatial transition from malignant cores to stromal boundaries. Spatial neighbors for each spot were defined by a *k*-nearest-neighbor graph (*k* = 6) constructed on the 10x Visium hexagonal lattice. Seed spots were assigned an initial layer index of Ls = 0, and all remaining spots were initialized as unassigned. At iteration *i*, each unclassified spot *s* was labeled according to the majority status of its spatial neighbors: spots with a malignant-dominated neighborhood (*n*_mal_ > *n*_norm_) were assigned as malignant with a positive layer index (*L_s_* = *i*), whereas spots with a normal-dominated neighborhood (*n*_norm_ > *n*_mal_) were assigned as normal with a negative layer index (*L*_s_ = -*i*). Spots without a clear majority remained unassigned. The propagation procedure was iterated for up to 20 rounds and terminated locally upon reaching stromal barriers or when no further assignments were possible.

### Prioritization of candidate TCR-T therapy targets

To prioritize candidate TCR-T therapy targets, we established three criteria and derived the corresponding gene sets, ultimately identifying the final candidates by intersecting these gene sets:

i. Cell Cycle Exclude: We excluded 50 cell cycle-associated genes from the 407 CTGs, retaining a subset of 357 candidate genes.
ii. Normal Tissue Low: Using bulk RNA-seq data from the GTEx database, we identified 133 genes exhibiting minimal expression (median TPM < 3) across all non-testicular normal tissue types.
iii. Malignancy Specific: We evaluated the malignancy specificity of CTGs using single-cell transcriptomic data from the Discovery Cohort. For a given CTG within a specific cancer type, its expression level was employed as a classification feature to discriminate malignant cells from normal reference cells and tumor microenvironment (TME) cells within each sample, yielding two AUROC values (AUROC_ref_ and AUROC_TME_). Overall malignancy specificity was quantified by the median sample-level AUROCs, and a gene was defined as highly malignancy-specific if both median metrics exceeded 0.55. For cancer types with insufficient samples containing normal reference cells (i.e., HGSOC, NB, and SKCM), only the TME metric (AUROC_TME_ ≥ 0.55) was considered. This analytical step yielded a subset of 82 genes.

### Statistical analysis

If not specified, the two-sided Wilcoxon rank-sum test was used to evaluate the differences between the two groups, and Spearman’s correlation test was used to assess the correlations between two variables. Multiple hypothesis testing was corrected using the Benjamini–Hochberg method. All analyses and visualizations were performed using R (v4.2.3).

## Data availability

All data supporting the findings of this study are available within the article and its accompanying Supplementary Information. Additional data are available from the corresponding author upon reasonable request.

## Code availability

All R scripts used for data processing and figure generation are available on GitHub (https://github.com/chunyangyyyyyyy/PanCancerCTG).

## Acknowledgements

The research is supported by National Natural Science Foundation of China (Fund 32370715, 82330108), Science Fund Program for Distinguished Overseas Young Scholars of Shandong Province (2023HWYQ-015), Taishan Young Scholar of Shandong Province (tsqn202312020), and the Cheeloo Young Scholar Program of Shandong University. The content is solely the responsibility of the authors and does not necessarily represent the official views of sponsors.

## Author contributions

C.F. and K.L. conceived and designed the study. C.F. performed the formal analyses.

C.F. and K.L. drafted the manuscript. S.Y., L.L., X.J. and X.L. provided critical feedback during manuscript preparation and revision. All authors read and approved the final manuscript.

## Supplementary Information

**Supplementary Figures 1–9**.

**Supplementary Dataset: Tables S1-S7**.

**Table S1.** Data sources for transcriptomic (sc/snRNA-seq, spatial, and bulk), DNA methylation, and ChIP-seq datasets.

**Table S2.** List of 1,294 CTG candidate genes.

**Table S3.** List of CTGs identified per cancer type and cell cycle-associated CTGs. **Table S4.** DNA methylation metrics (MML and HSI) of CTGs across TCGA cancer types.

**Table S5.** Reactome pathway enrichment results for non-CTG genes co-expressed with CTG subgroups.

**Table S6.** Benchmarking of MaligCTG and established methods for malignant cell identification.

**Table S7.** Evaluation metrics and peptide predictions for prioritized candidate TCR-T targets.

