## Supplemental Figures for "Pan-cancer landscape of cancer-testis genes revealed by single-cell and spatial transcriptomics"

Chunyang Fu *et al.*

**This PDF file includes:**

Supplementary Figures 1–9

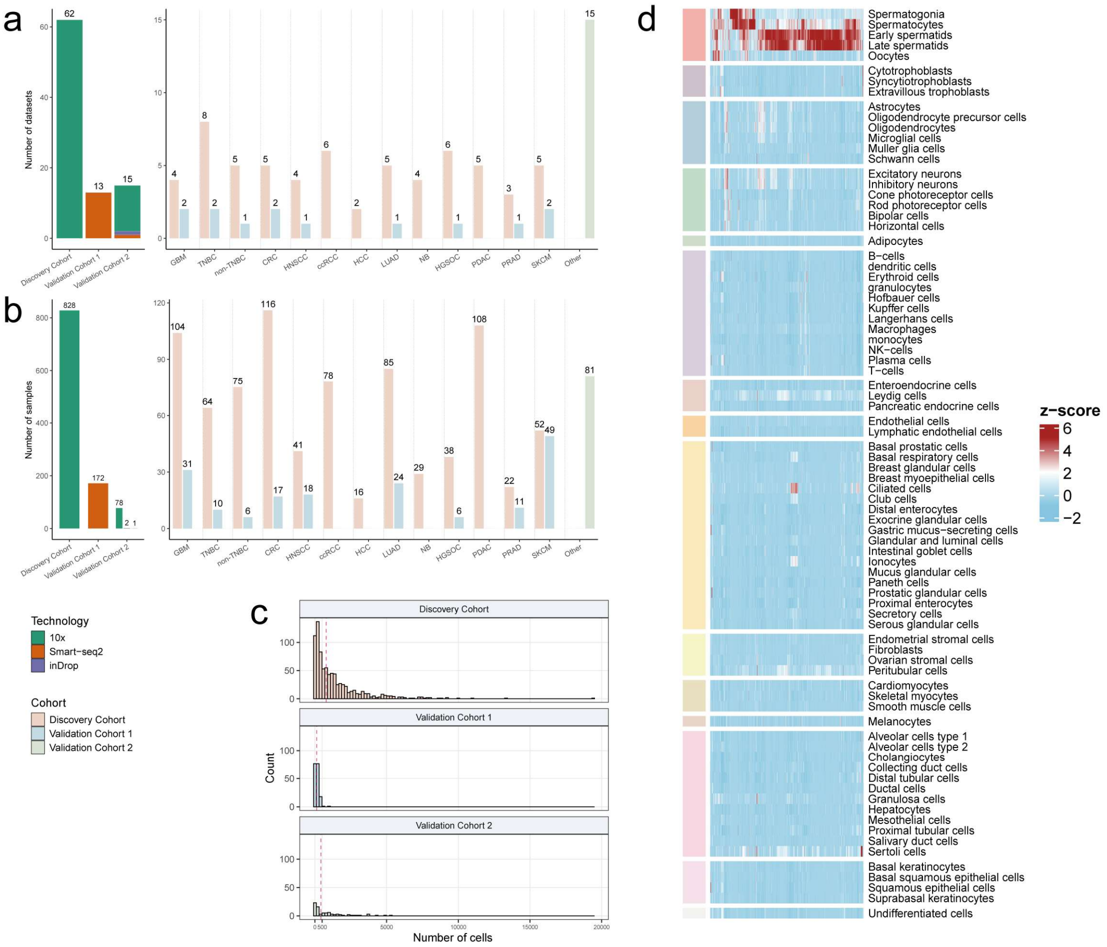


**Supplementary Fig. 1. Overview of curated sc/snRNA-seq datasets and candidate genes**.

**(a)** Bar plots showing the dataset count per cohort, stratified by sequencing platform (left), and the dataset count per cancer type, stratified by cohort (right).

**(b)** Bar plots showing the sample count per cohort, stratified by sequencing platform (left), and the sample count per cancer type, stratified by cohort (right).

**(c)** Histograms of malignant cell counts per sample across the three cohorts; red dashed line denotes the median.

**(d)** Heatmap of column-scaled expression for candidate genes across 81 normal cell types. RNA expression data was obtained from HPA (https://www.proteinatlas.org/).

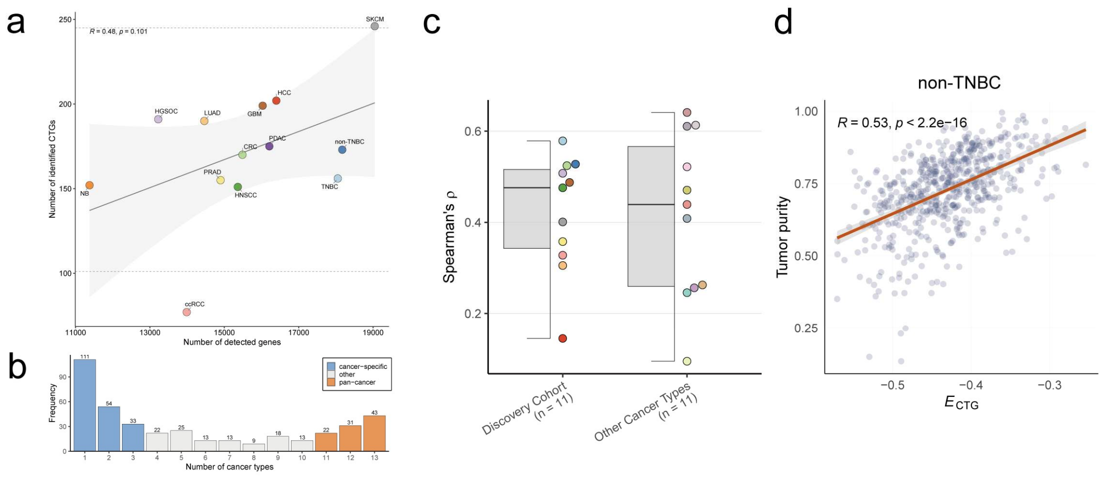


**Supplementary Fig. 2. Characteristics of the identified CTGs.**

**(a)** Scatter plot illustrating the relationship between the number of detected genes (x-axis) and the number of identified CTGs (y-axis) across cancer types. The solid grey line represents a linear regression fit, with the shaded region indicating the 95% confidence interval. Spearman correlation and *P*-value are shown. Horizontal dashed lines indicate the outlier boundaries defined by the interquartile range (IQR) method (Q1 − 1.5 × IQR and Q3 + 1.5 × IQR).

**(b)** Bar plot depicting the frequency distribution of CTGs based on the number of cancer types in which they were identified.

**(c)** Boxplot showing the Spearman correlation coefficients between aggregate CTG expression and tumor purity for cancer types in the Discovery Cohort and other TCGA cancer types. Each point represents a cancer type.

**(d)** Scatterplot illustrating the correlation between aggregate CTG expression and tumor purity specifically in non-TNBC tumor samples from TCGA. Each point represents an individual tumor sample.

Boxplots represent the median (center line), upper and lower quartiles (box limits), and 1.5 × IQR (whiskers).

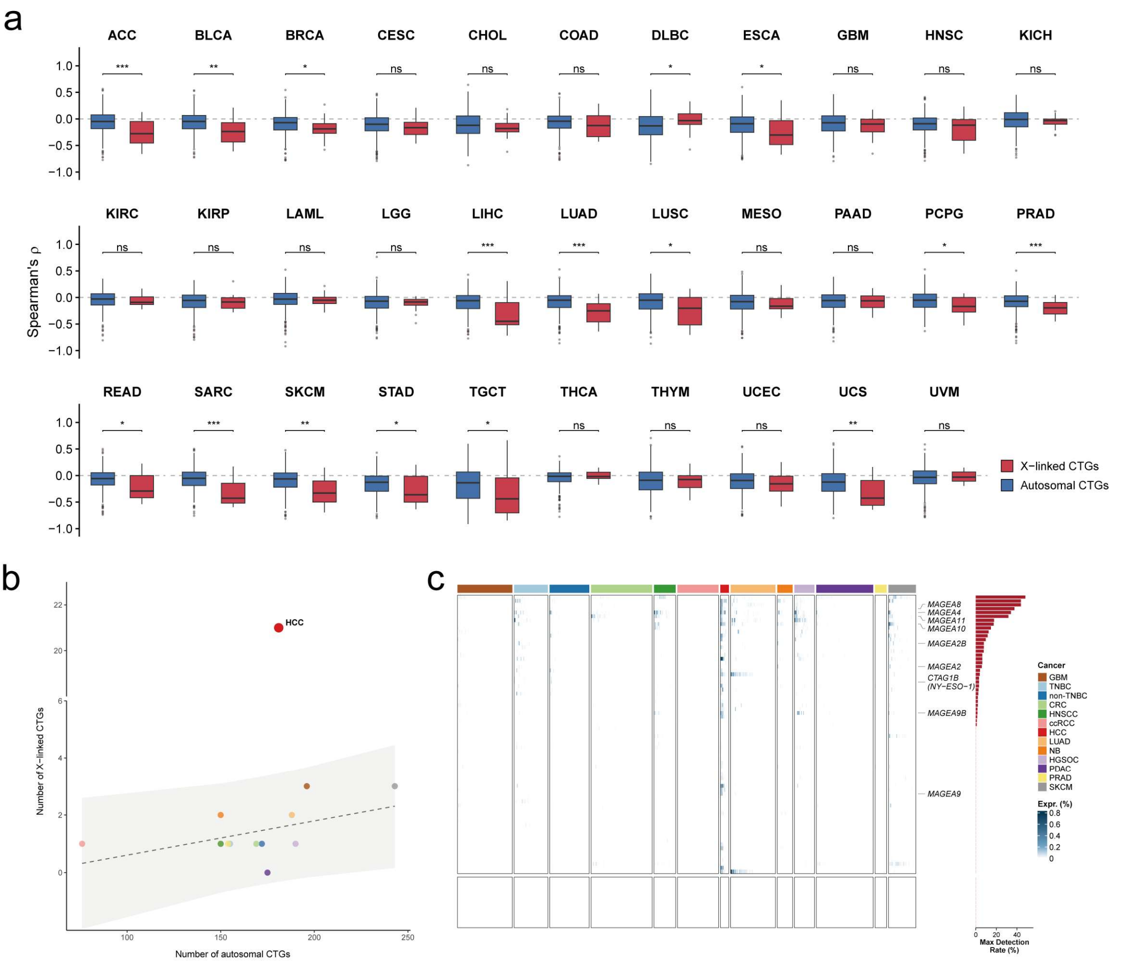


**Supplementary Fig. 3. DNA methylation dependency dictates the sample-restricted expression of X-linked CTGs.**

**(a)** Boxplots illustrating the Spearman correlation coefficients between gene expression and promoter methylation for X-linked and autosomal CTGs across TCGA cancer types.

**(b)** Scatter plot illustrating the relationship between the number of identified autosomal CTGs and X-linked CTGs across different cancer types. The linear regression model (dashed line) and 95% prediction interval (grey shaded area) were generated using data from all cancer types excluding HCC.

(**c**) Heatmap displaying the percentage of malignant cells expressing the X-linked CTGs (recorded in CTdatabase but not included in our CTG set) across tumor samples. The right-sided bar plot represents the maximum detection rate for each gene across all evaluated cancer types. For visualization purposes, missing data are rendered as zeros (white) in the heatmap only.

Boxplots represent the median (center line), upper and lower quartiles (box limits), and 1.5 × IQR (whiskers). Significance: $^{*}P \leq 0.05$, $^{**}P \leq 0.01$, $^{***}P \leq 0.001$, $^{****}P \leq 0.0001$; ns, not significant.

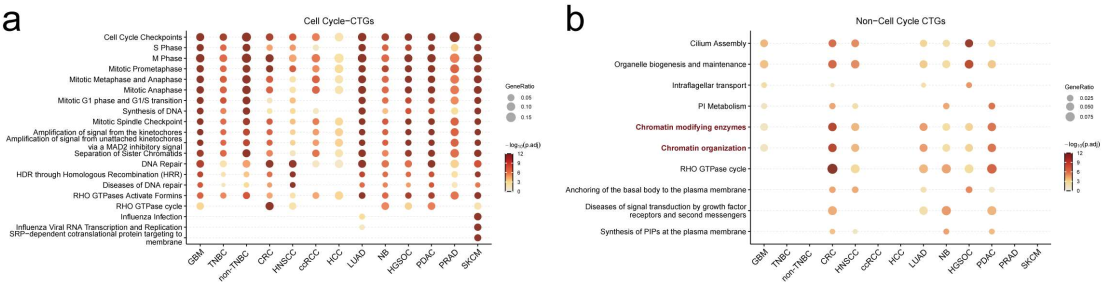


**Supplementary Fig. 4. Pathway enrichment analysis of non-CTG genes co-expressed with distinct CTG subgroups.**

**(a–b)** Reactome pathway enrichment analysis of the top 2,000 non-CTG genes exhibiting the highest frequency of significant positive co-expression with **(a)** cell cycle-associated and **(b)** non-cell cycle-associated CTGs across the Discovery Cohort.

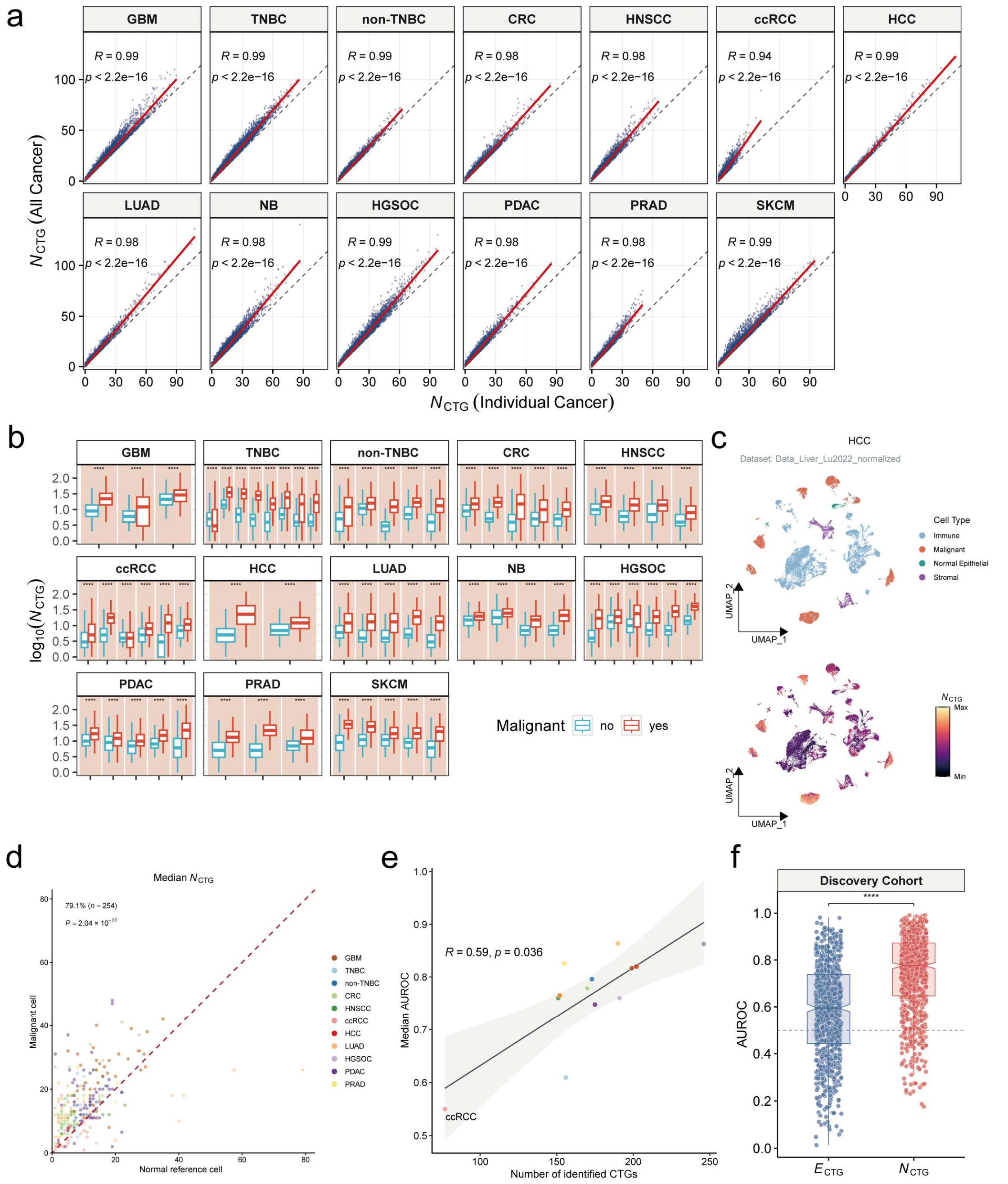


**Supplementary Fig. 5. Robust elevation of $N_{CTG}$ in malignant cells across sc/snRNA-seq datasets**.

**(a)** Scatterplots correlating $N_{CTG}$ values computed using all 407 CTGs (y-axis) versus only the CTGs identified in the specific cancer type (x-axis). For visualization, 1,500 cells were randomly sampled per dataset from the Discovery Cohort. The dashed grey line indicates $x = y$, and the red solid line represents the linear regression fit. Spearman's correlations and $P$-values are shown.

**(b)** Boxplots comparing $N_{CTG}$ in malignant versus non-malignant cells for each dataset in the Discovery Cohort.

**(c)** UMAP plots visualizing cell annotations (top) and $N_{CTG}$ levels (bottom) in a representative dataset from the Discovery Cohort.

**(d)** Pairwise comparisons of the median $N_{CTG}$ between malignant cells (y-axis) and corresponding normal reference cells (x-axis) within the same samples from the Discovery Cohort. Each dot represents an individual tumor sample and is color-coded by cancer type. The red dashed line represents the line of $y = x$. The percentages indicate the proportion of samples wherein malignant cells harbor a higher median $N_{CTG}$ than normal reference cells.

**(e)** Scatterplot illustrating the relationship between the number of identified CTGs and the median AUROC values, which quantified the separation between malignant and non-malignant cells using $N_{CTG}$ across samples in a given cancer type. Spearman's correlations and *P*-values are shown.

**(f)** Boxplots of AUROC values from the Discovery Cohort using different indices: aggregate CTG expression ($E_{CTG}$) and $N_{CTG}$.

*P*-values were determined using two-sided paired Wilcoxon signed-rank tests for comparisons in panels **(d)** and **(f)**. Boxplots represent the median (center line), upper and lower quartiles (box limits), and 1.5 × IQR (whiskers). Significance: $*P \leq 0.05$, $**P \leq 0.01$, $***P \leq 0.001$, $****P \leq 0.0001$; ns, not significant.

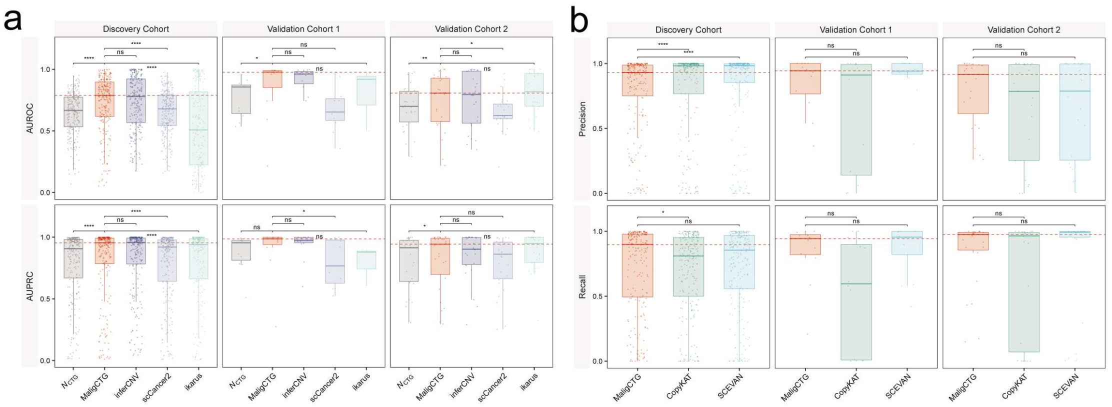


**Supplementary Fig. 6. Evaluation of MaligCTG performance on malignant and normal reference cell populations.**

**(a)** Boxplots comparing the continuous scoring accuracy (AUROC and AUPRC) of MaligCTG against $N_{\mathrm{CTG}}$, inferCNV, scCancer2, and ikarus across three cohorts.

**(b)** Boxplots benchmarking the binary classification performance (precision and recall) of MaligCTG against CopyKAT and SCEVAN across three cohorts.

In contrast to **Fig. 4**, the analyses in **(a)** and **(b)** were restricted to malignant cells and normal reference cells. *P*-values were determined using two-sided paired Wilcoxon signed-rank tests. Boxplots represent the median (center line), upper and lower quartiles (box limits), and 1.5 × IQR (whiskers). Points represent individual tumor samples. Significance: $*P \leq 0.05$, $**P \leq 0.01$, $***P \leq 0.001$, $****P \leq 0.0001$; ns, not significant.

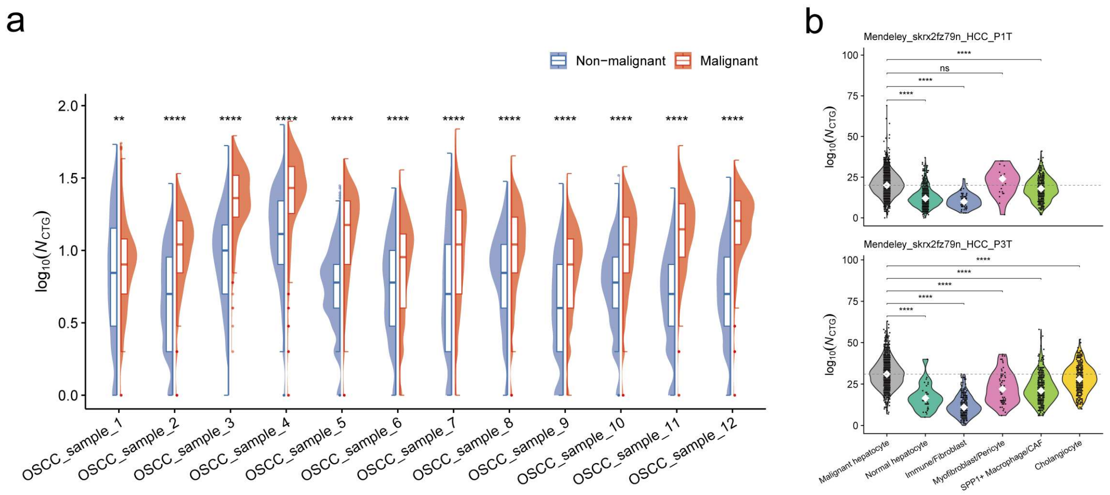


**Supplementary Fig. 7. Elevation of $N_{\mathrm{CTG}}$ in malignant spots**.

**(a)** Violin plots comparing $N_{\mathrm{CTG}}$ between histologically annotated malignant and non-malignant spots in 12 OSCC samples.

**(b)** Violin plots of $N_{\mathrm{CTG}}$ in malignant versus normal cell types for two HCC samples.

*P*-values were determined using two-sided Wilcoxon rank-sum tests for comparisons in all panels. Significance: $*P \leq 0.05$, $**P \leq 0.01$, $***P \leq 0.001$, $****P \leq 0.0001$; ns, not significant.

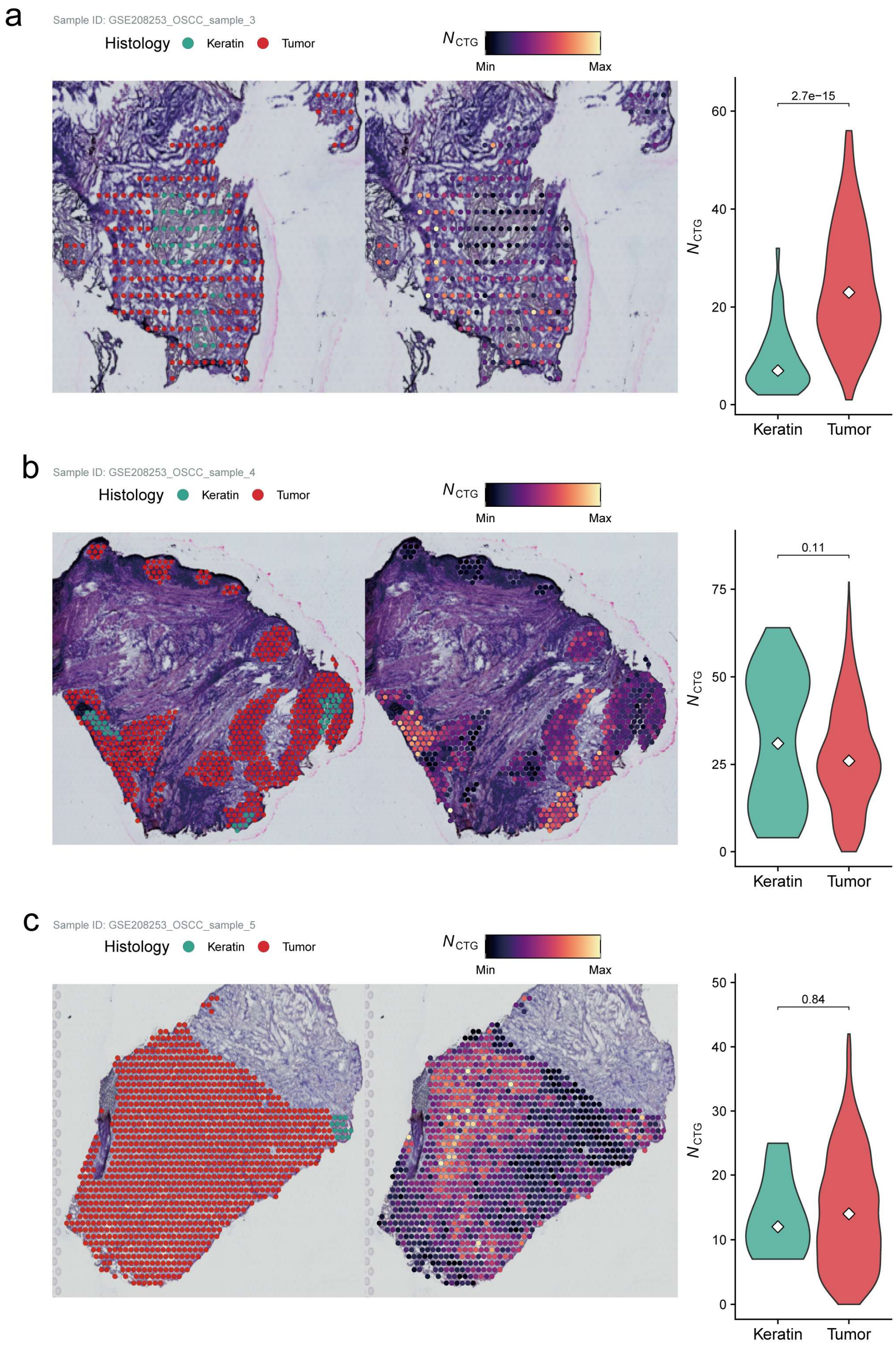


**Supplementary Fig. 8. Comparison of $N_{CTG}$ levels between keratin and tumor regions.**

**(a–c)** Spatial visualization of pathologist-annotated keratin and tumor regions (left) and $N_{CTG}$ values in these regions (middle) in three additional HNSCC samples. Accompanying violin plots (right) quantify the differences in $N_{CTG}$ levels between tumor and keratin regions for each sample.

*P*-values were determined using two-sided Wilcoxon rank-sum tests for comparisons in all panels. Significance: $^{*}P \leq 0.05$, $^{**}P \leq 0.01$, $^{***}P \leq 0.001$, $^{****}P \leq 0.0001$; ns, not significant.

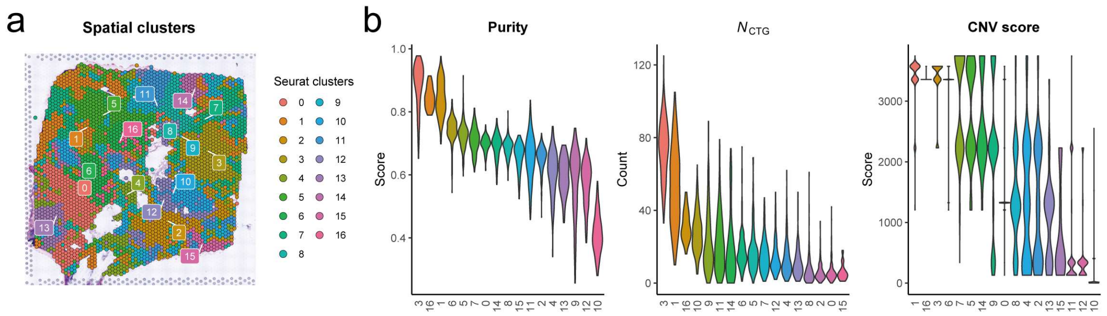


**Supplementary Fig. 9. Spatial clustering of a representative breast cancer sample.**

**(a)** Spatial visualization of the Seurat clustering results (resolution = 1.5) for the representative BRCA sample (Sample ID: HTAN_HT206B1-U1_ST_Bn1) shown in **Fig. 6c**.

**(b)** Violin plots depicting the tumor purity (left), $N_{\mathrm{CTG}}$ (middle), and CNV score (right) across Seurat clusters in this sample.
